# Automated 3D Inner Ear Assessment for Optimizing Animal Models of Noise-Induced Hearing Loss

**DOI:** 10.1101/2020.09.30.317941

**Authors:** Lei Shi, Yi Du, Cong Wei, Wen-jie Huang, Han-dai Qin, Chen Liu, Da Liu, Shuo-long Yuan, Qing-qing Jiang, Jun Yang, Gang Li, Hong-lan Xie, Jian-guo Zhao, Ning Yu, Shi-ming Yang

## Abstract

**Objective:** To facilitate the selection of optimal animal models for noise-induced hearing loss (NIHL) research, this study aims to develop an accurate, automatic centerline-based method to construct three-dimensional (3D) inner ear models in various mammals and assess their morphological similarities to humans.

**Methods:** Three adult mice, three guinea pigs, three mini pigs, and one human temporal bone were scanned using Micro-CT. After segmentation, 3D inner ear models were reconstructed. A novel centerline method was established to automatically estimate 3D properties and calculate length, volume, and angle parameters across species.

**Results:** Detailed 3D models of the cochlea, vestibule, and semicircular canals were successfully constructed. Mean lengths and volumes of the cochlea and semicircular canals generally increased with body size, with mini pigs demonstrating the closest proximity to human data. Notably, numerical assessments showed that the semicircular canals in both mice and mini pigs closely resembled human structures.

**Conclusion:** Automatic centerline-based measurement provides an effective tool for assessing 3D inner ear morphology. By confirming the high morphological similarity between mini pigs and humans, this study provides a critical theoretical basis for the mechanical analysis of noise-induced damage and the selection of biomimetic animal models for NIHL studies.

## 1. Background

The osseous labyrinth, located in the petrous portion of the temporal bone, consists of the vestibule, semicircular canals, and cochlea. All these small and irregular inner ear components and their interspace relationships are vital in surgical procedures and basic studies on auditory mechanics ^[1–3]^. Precise geometric characterization of the inner ear is essential for understanding its physiological functions. To this end, various imaging techniques have been employed to capture topographic data, including traditional histology, computed tomography (CT), magnetic resonance imaging (MRI), and synchrotron radiation-based micro-CT ^[1, 4–8]^. Advanced post-processing and 3D visualization enable the precise quantification of inner ear structures. These rotatable 3D models facilitate a comprehensive understanding of anatomical features and structural variations, while providing a robust framework to evaluate the morphological similarity between various animal models and the human inner ear.

This, in turn, allows for the selection of more reliable animal models for noise-induced hearing loss (NIHL) research ^[3, 8, 9]^. Considering density resolution and operation procedures, CT seems to be the optimal method to obtain inner ear topography ^[3, 4, 8–10]^. Micro-CT is a variation of CT with a higher resolution and could meet the requirement of remodeling fine inner ear structures ^[4, 11]^.

Mammalian models are indispensable in otology research for visualizing temporal bone anatomy, evaluating surgical trauma, and establishing NIHL models ^[1–6, 8–10, 12–15]^. Accurate quantification of inner ear dimensions is crucial for mechanical modeling and analyzing acoustic stress distribution^[16, 29]^. A comprehensive characterization of the morphological features of these animal models is essential for advancing biomechanical insights into the screening of NIHL models ^[11, 14, 17]^. However, quantitative analysis of these different 3D models depends highly on elaborate preparation, image resolution, and iterative measurements, which may cause radiation damage or inevitable error and eventually affect the accuracy and validity of results ^[1, 8, 9]^. Furthermore, the lack of comparative quantitative data between mammals limits the scientific selection of optimal animal models for human NIHL research.

Although the inner ear structure is complicated, the cochlear is very close to a spiral shape, and the semicircular canals are close to circles. Recently, the centerline method, successfully used in the segmentation of the vessel^[18]^, vocal tract^[19, 20]^, and cochlear^[12]^, shows good properties in path tracking, which make it possible to describe the feature of intricate solid objects like inner ear.

Accordingly, this study employs the centerline method to overcome the limitations of traditional 3D imaging, enabling the automatic extraction of morphological parameters across different mammals. This advanced approach facilitates structural research and provides a mathematical basis for identifying the most suitable animal proxy for human NIHL. By comparing various models with human data, we describe the discrepancies in the labyrinthine structure mathematically.

## 2 Materials and Methods

### 2.1 Preparation of mammal specimens

To investigate the differences between animals, healthy adult mice, guinea pigs, and mini pigs (n=3 in each group) were included in this research. A left human temporal bone without any evidence of malformation was analyzed to compare the variations between animals and humans.

For getting an image of an intact inner ear, the animal samples were prepared as follows. After an intraperitoneal injection of a lethal dose of sodium pentobarbital, mice, guinea pigs, and mini pigs were sacrificed. Following decapitation, the heads of the mice and the guinea pigs were divided, and both their left and right bullaes were removed for image acquisition. The left and right temporal bones of mini pigs were cut right after decapitation, including the complete middle and inner ear. All the specimens were immediately scanned after the above steps. Iterative procedures for sample preparation, unavoidably leading to shrinkage in the samples ^[8]^, are not necessary to our study, which primarily aimed at testing the efficiency of a novel measuring method. Therefore, no steps of fixation, dehydration, and decalcification were processed in our research.

### 2.2 Micro-CT scanning procedure

All the prepared samples were scanned using a high-speed Micro-CT system (PerkinElmer Inc., Norwalk, CT, USA). A 5×5mm view was used in the samples of mice, a 20×20mm field of view in guinea pigs and mini pigs, 30×30mm view in the human temporal bone. The scan was performed at a voltage of 90 kV and a current of 120 μA. The fast scan time of 4.5min was set, and 512 slices of the image with a voxel size of 9μm, 39μm, and 59μm were achieved in these cases. The data set was exported to disc in the type of DICOM after the process of the workstation. No defects or infections were shown in all three samples.

### 2.3 3D reconstruction

Then MIMICS (Materialises interactive medical image control system, Belgium) was used in this research to process the images and reconstruct 3D volume. The function of interpolation in the MIMICS allows a semi-automated segment between the slices ^[4]^. Firstly, the segmentation was achieved automatically using different gray values, separating the object from the background. To remove artifacts not belong to the object, manual segmentation was supplemented to fit the target region. The segmentation results were checked and adjusted in transversal and coronal planes as well as 3D views.

After defining the mask in involving slices, the region of interest was determined. Then calculate the mask, 3D volume of the labyrinth embedded in the temporal bone was rendered (Fig.1). Further smooth and modification were carried on for calculation.

**Fig. 1:**
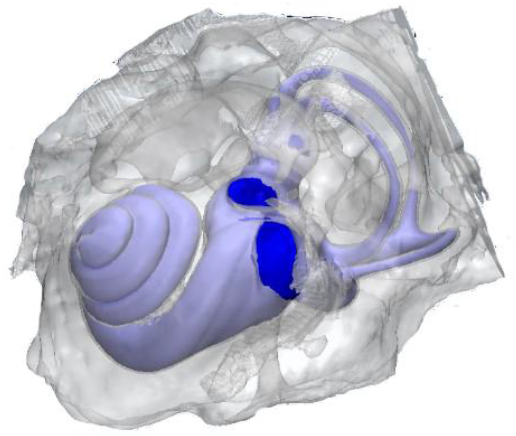
Volume render of the left temporal bone and inner ear in mini pig model. The temporal bone is rendered translucent.

### 2.4 Automatic Measurements of 3D models

#### 2.4.1 Calculate Centerline

Fit centerline, one of the Medcad modules in Mimics, was used to automatically calculate the centerlines of 3D models. After finishing centerline calculation, control points could be easily added if necessary. The centerlines and nodes of models were illustrated in Fig.2 and Fig.3.

**Fig. 2:**
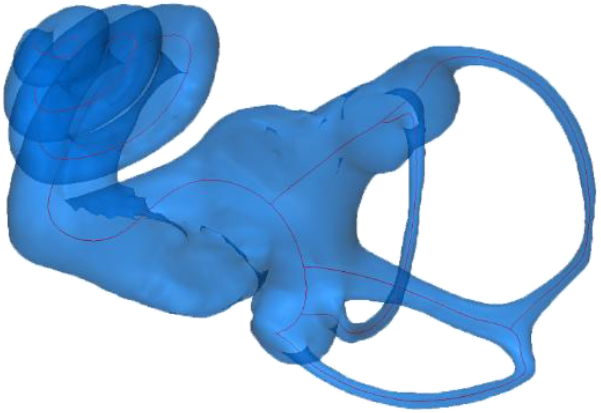
Centerlines in 3D model of inner ear in mini pig

**Fig. 3:**
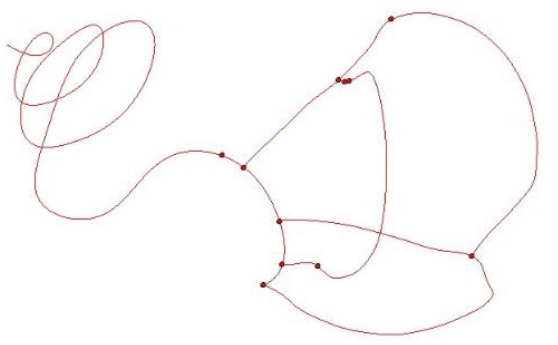
Centerlines and nodes of inner ear in mini pig

#### 2.4.2 Automatic measurements of inner ear parameters in various samples

The unique nodes in the centerlines were used as landmarks. The landmarks in the 3D models were mapped to corresponding anatomic locations in inner ear. The distances between the landmarks were used to calculate the length automatically. The length of the cochlea was automatically measured from the landmark of apex of the cochlea to the round window niche (fiburcation) through the centerline. The length of anterior and posterior semicircular canal were measured from the feature points representing ampulla to common crus. The length of lateral semicircular canal was measured between two foot points. The common crus length was also measured by two foot points.

Based on the landmarks, the 3D model of the labyrinth was segmented into six parts, cochlea (CO), vestibule (VL), anterior semicircular canal (ASC), lateral semicircular canal (LSC), posterior semicircular canal (PSC) and common crus (CC). Then, the volumes of these six components were computed automatically in Mimics software.

The centerlines was exported in the form of IGS and imported into a geometry reconstruction software Geomagic Studio (Geomagic, Morrisville, North Carolina, USA) for further analysis. Reference plane in corresponding semicircular canals was defined with the use of plane of best fit in feature modules (shown in Fig.4). Take the anterior SC and lateral SC as an example, shown in Fig. 5, the method of obtaining the plane included angle is as follows: 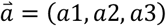 is the vector of the anterior SC reference plane; 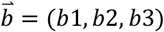 is the vector of the lateral SC reference plane; the angle between the two intersection plane is described in Eq.(1).

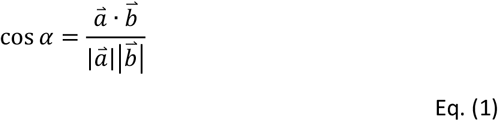

**Fig. 4:**
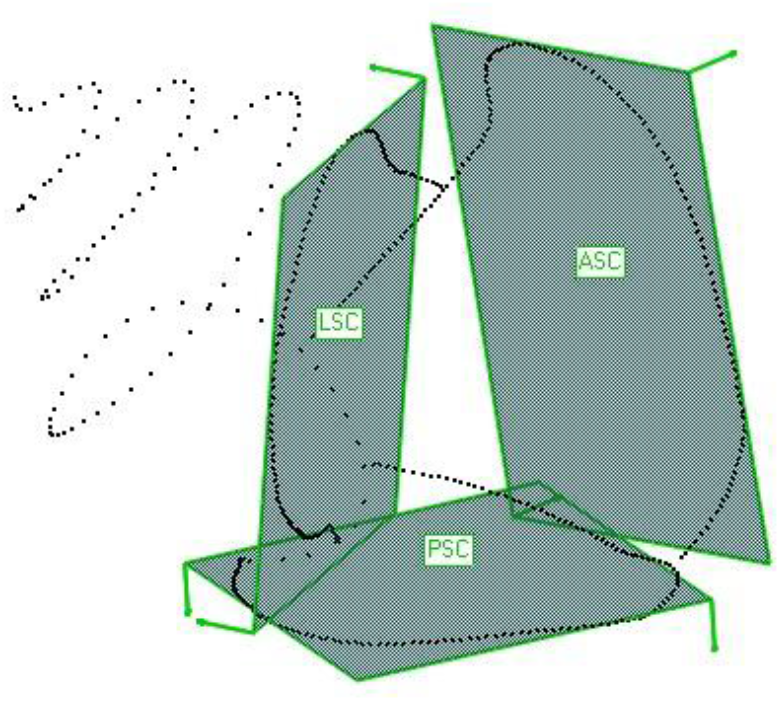
Illustration of the best fit plane in mini pig model, ASC is the reference plane of anterior semicircular canal, LSC is the reference plane of lateral semicircular canal, PSC is the reference plane of posterior semicircular canal

**Fig. 5:**
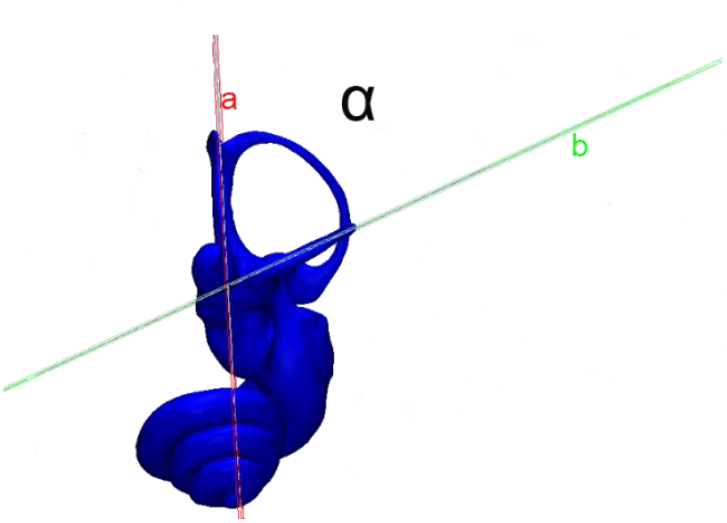
Illustration of the angle measurement in mini pig model, a: axis of the plane of the anterior semicircular canal; b: axis of the plane of the lateral semicircular canal; α: included angle between a and b

The algorithm to calculate the angles was realized through Matlab platform (MATHWORKS INC, Natic, USA).

## 3 Results

### 3.1 Geometric reconstruction of the inner ear renders exquisite structures of CO, VL, and SC

The intricate, elaborate structure of the inner ear model was rendered as a complex geometric solid comprising of the cochlea, vestibule, and semicircular canals, shown in Fig. 6. The 3D visualization made it possible to observe the surface texture and anatomical details of the inner ear, such as the apical and basal turn, and the round window and oval window. The spatial location and concrete form of the semicircular canals are embodied in 3D models. Three similar-circular canals owned similar lengths and were visually perpendicular to each other. Common crus and ampulla were also easy to recognize from the models. To our knowledge, the detailed 3D visualization of the mini pig inner ear in our research is shown for the first time.

**Fig. 6:**
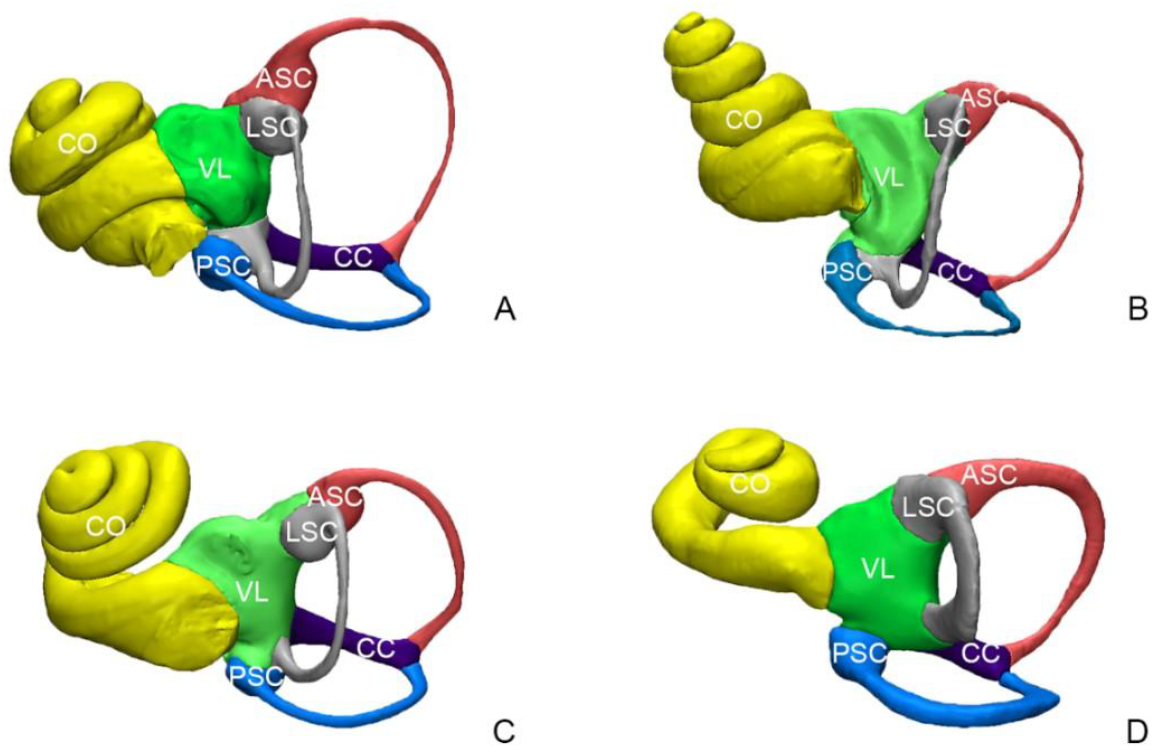
Illustration of inner ear structure of different mammals.(A: mouse; B: guinea pig; C: mini pig; D: human) CO-cochlea; VL-vestibule; LSC-lateral semicircular canal; ASC-anterior semicircular canal; PSC-posterior semicircular canal; CC-common crus

### 3.2 The length parameters of the inner ear increase with the body size of mammals

Fig. 7 showed the morphology of centerlines and landmarks in three animal models and the human model. The centerlines and landmarks formed a framework of a brief characteristic description of inner ears. Characteristics in centerlines mapped to the morphological features of three-dimensional models, such as oval window, ampulla, and common crus. The cochlear turns varied among animals, the specifical mouse is two turns, the guinea pig is four turns and 3.5 turns for the mini pig, but the human cochlear is less than three turns(Fig.8).

**Fig. 7:**
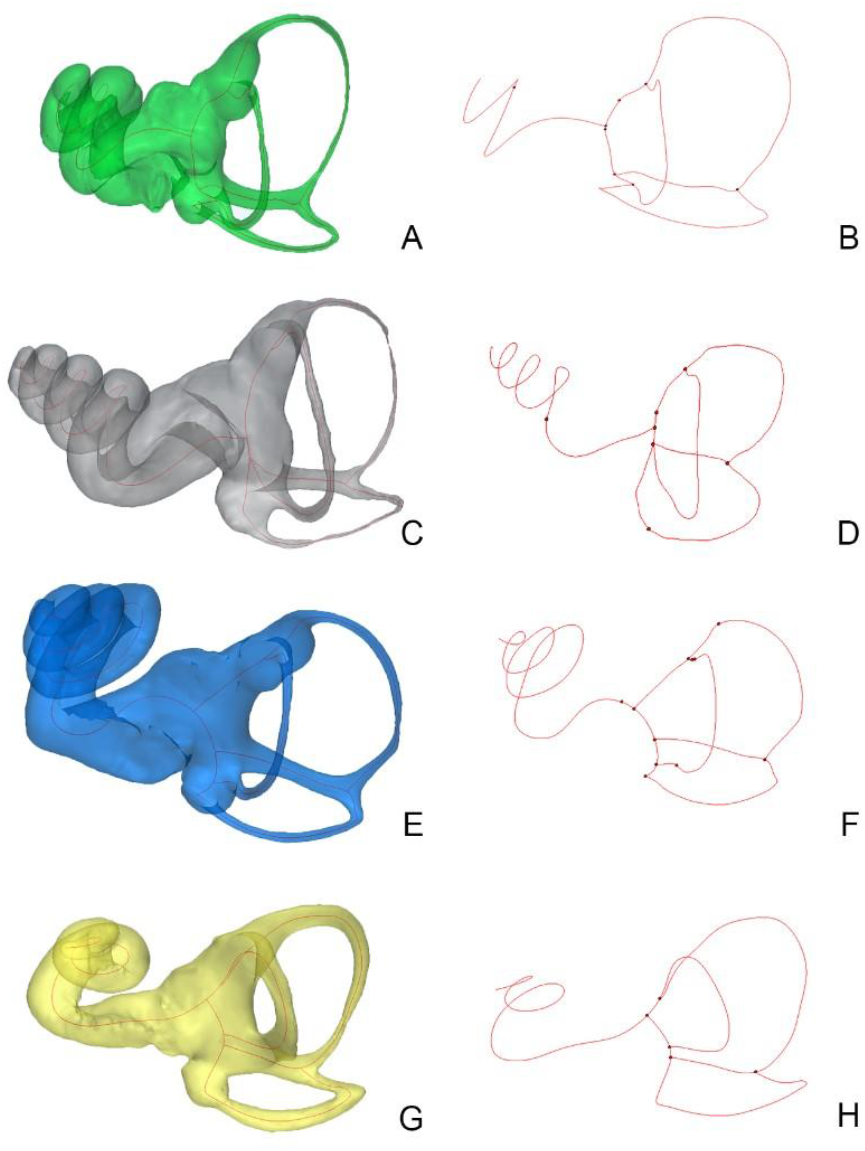
Centerlines and landmarks in various mammal models (A,B: mouse inner ear model(A) and its centerlines and landmarks(B); C,D: guinea pig inner ear model(C) and its centerlines and landmarks(D); E,F: mini pig inner ear model(E) and its centerline and landmarks(F); G,H: human inner ear model(G) and its centerline and landmarks(H)

**Fig. 8:**
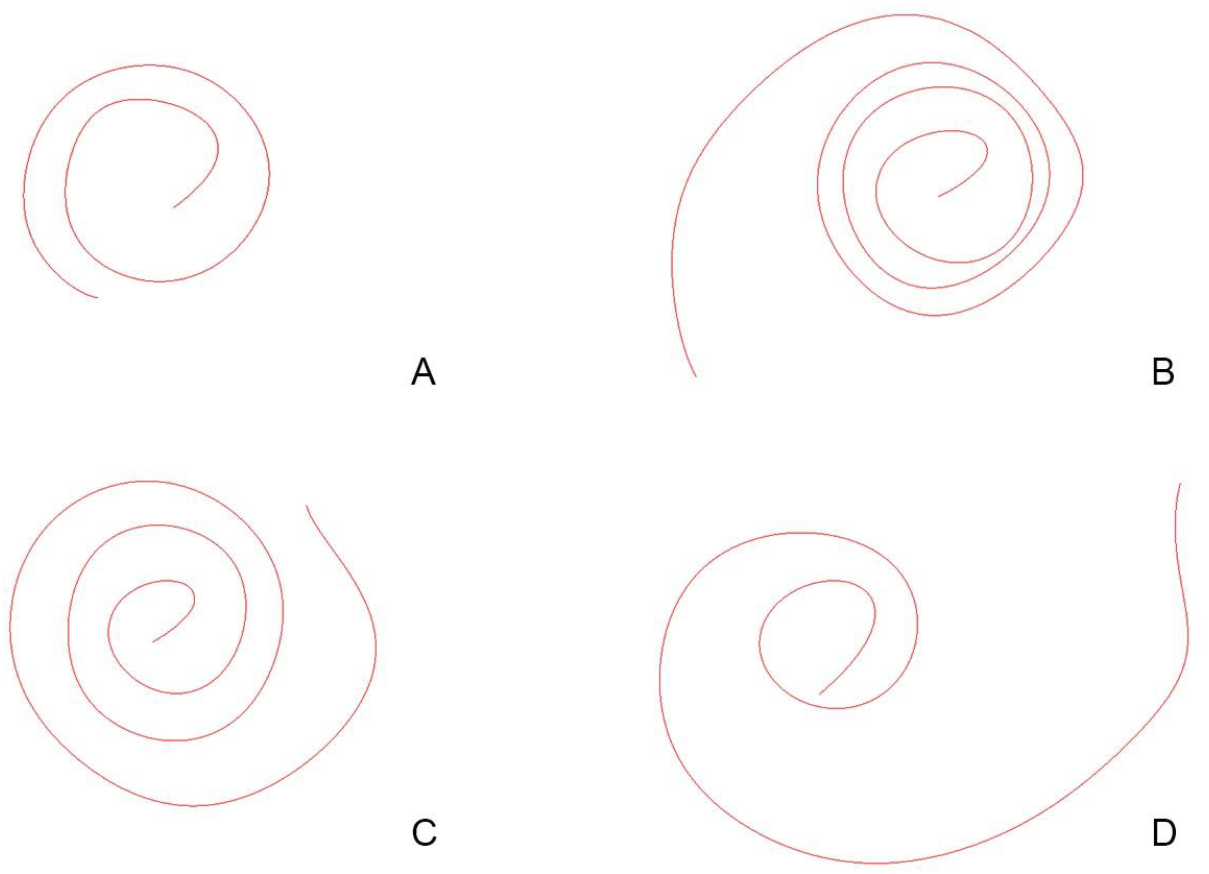
Cochlear turns of different mammals (A: mouse cochlear; B: guinea pig cochlear; C: pig cochlear; D: human cochlear)

The length measurements of inner ear models were listed in Table 1, including the cochlea, lateral semicircular canal, anterior semicircular canal, posterior semicircular canal, and common crus. Among all the measurements, the cochlear length was the maximum distance. The lengths of three semicircular canals were similar to each other. Lengths increased with body size, and among them, the mean value of the cochlear length of mini pigs was the closest to humans.

**Table 1.** Length Measurements of Inner Ear.

| Length<br>(mm) | CO | LSC | ASC | PSC | CC |
| --- | --- | --- | --- | --- | --- |
| Mice | 6.14±0.36 | 3.54±0.35 | 4.28±0.33 | 3.90±0.35 | 1.33±0.13 |
| Guinea pigs | 17.65±0.53 | 8.14±0.36 | 7.60±0.28 | 7.52±0.69 | 2.80±0.09 |
| Mini pigs | 26.09±1.22 | 9.72±0.33 | 9.87±0.67 | 9.36±0.67 | 3.98±0.34 |
| Human | 24.95 | 13.26 | 16.24 | 16.93 | 4.15 |

The value represents mean±SD for the group of mice, guinea pigs and mini pigs

### 3.3 The volume parameters of the inner ear increase with the body size of animals

Fig. 6 illustrated the 3D model of different animal inner ears with segmented and color-marked components: cochlea (yellow), the vestibule (green), the anterior semicircular canal (red), the lateral semicircular canal (gray), the posterior semicircular canal (blue) and common crus (purple), the volume of each part was calculated automatically in Mimics. Generally, the inner ear volume of mice, Guinea pigs, and mini pigs was less than that of humans and increased with the body size. In each experimental animal, the most considerable portion was cochlea; the next is a vestibule, three semicircular canals, and the least is common crus. Detailed data was shown in Table 2.

**Table 2.** Volume Measurements of Inner Ear.

| Volume<br>(mm<sup>3</sup>) | CO | VL | LSC | ASC | PSC | CC |
| --- | --- | --- | --- | --- | --- | --- |
| Mice | 1.54 ± 0.27 | 0.55 ± 0.05 | 0.18 ± 0.05 | 0.15 ± 0.01 | 0.14 ± 0.03 | 0.07 ± 0.01 |
| Guinea pigs | 13.68 ± 1.51 | 7.11 ± 0.89 | 1.52 ± 0.29 | 0.79 ± 0.14 | 0.80 ± 0.20 | 0.63 ± 0.26 |
| Mini pigs | 40.97 ± 5.41 | 20.53 ± 2.13 | 3.13 ± 0.92 | 2.78 ± 0.66 | 2.41 ± 0.69 | 1.65 ± 0.30 |
| Human | 59.37 | 46.65 | 13.70 | 9.85 | 12.25 | 4.28 |

The value represents mean ± SD for the group of mice, guinea pigs and mini pigs

### 3.4 The angle parameters of mini pig inner ear are closest to the human inner ear

The spatial relationship between the semicircular canals was illustrated in Table 3(the mice’s angle parameters, guinea pigs, and mini pigs were shown in Mean±SD). The three reference planes are mutually perpendicular in vision, whereas the angle parameter differed from each other. Our research also compared the angle discrepancy between experimental animal models and human data with the use of Euclidean distance. The detailed distance was calculated by Eq. (2).

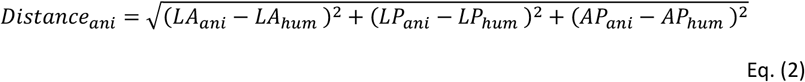

**Table 3.** Angle Measurements between the Semicircular Canals.

|  | L-A SC<br>(°) | L-P SC<br>(°) | A-P SC<br>(°) | Euclidean<br>distance |
| --- | --- | --- | --- | --- |
| Mice | 95.84±0.33 | 97.46±0.80 | 91.54±2.31 | 16.10 |
| Guinea pigs | 100.77±4.18 | 83.93±1.88 | 87.43±2.43 | 23.13 |
| mini pigs | 71.37±4.01 | 83.45±1.07 | 86.15±2.15 | 16.33 |
| Human | 81.40 | 96.24 | 84.53 | 0 |

Abbreviations: L-A SC = angles between lateral and anterior semicircular canals; L-P SC = angles between lateral and posterior semicircular canals; A-P SC = angles between anterior and posterior semicircular canals. The value represents mean±SD for the group of mice, guinea pigs and mini pigs.

The *Distance*_ani_ is the Euclidean distance between each animal and human, LA is the angle between lateral and anterior semicircular canal reference planes, LP is the angle between lateral and posterior semicircular canal reference planes, AP is the angle between anterior and posterior semicircular canal reference planes, ani represents angle parameter in each animal (mice, guinea pigs, and mini pigs) and hum is the angle data in human. The results of Euclidean distance indicated that the angle parameters of mice are closest to human in measurements, the mini pigs are also very similar, the least are guinea pigs; namely, the spatial relationship of semicircular canals was more approximate to human in mice and minipigs than that in guinea pigs.

## 4. Discussion

This study characterizes the morphological disparities of the inner ear across various mammalian models, providing the first 3D visualization and precise quantification of the minipig inner ear to assess its potential for hearing loss research. Crucially, all dimensional, volumetric, and angular parameters of the minipig inner ear exhibit a higher degree of similarity to human values than those of mice or guinea pigs—species traditionally utilized in NIHL studies. While the anatomical similarity of the minipig middle ear to humans has been established ^[21, 22]^, our findings extend this by providing a high-precision evaluation of the cochlear micro-structure. Due to different measuring methods and animal samples, the above literature had described that the length of mini pig cochlear was approximately 39.48mm coiling three and a half times, which was larger than our current research(mean value is 26.09mm). The mean length of mice cochlear in our measurement was 6.14mm, approaching previous data ranging from 5.81mm to 6.67mm ^[5, 23, 24]^. Our data on mice cochlear volume of 1.54mm^3^ resembled Buytaert’s result (1.702mm^3^) ^[8]^. The guinea pig cochlear, the mean length was 17.65mm, which was consistent with other researches ranging from 15.7mm to 22mm ^[14, 25]^. Different imaging acquisition and measuring methods, volume, and length of the human inner ear were slightly underestimated in our research compared to previous data ^[1, 4, 7, 9, 26]^. Melhem et al. reported the female human inner ear’s volumetric measurements to range from 172.3mm^3^ to 241.9mm^3^, indicating that there are some variabilities in the length and volume of the inner ear between individuals ^[7]^.

The semicircular canals are sensory organs that are sensitive to angular accelerations of the head ^[27]^. Understanding the planer orientation of semicircular canals is important to investigate compensatory eye movements due to the vestibule-ocular reflex, which occurs parallel to each semicircular canal plane, the basis for the test and treatment for the dizzy patient^[27, 28]^. Although three semicircular canals were perpendicular to each other visually, the plane included angles that were different in precise measurements. With the method of a histologic section, Hashimoto et al. reconstructed the 3D models of semicircular canals of 7 human temporal bones and reported the average angle for each pair of semicircular canals is approximately 90°^[27]^. Aoki et al. investigated 11 persons with intact vestibular function with MR scan and reported that the average of L-A SC is 91.6°, L-P SC is 90.8°, and A-P SC is 96.2° ^[28]^. Christensen et al. studied ten human data sets and calculated that the average of L-A SC is 92.79°, L-P SC is 90.93°, and A-P SC is 100.75° ^[1]^. Our result of the human model shows a little difference with previous data due to different measuring methods. Former angle measurements were commonly based on manual process ^[1, 28]^ and complicated equations ^[7]^ that would cause an inevitable error and influence data accuracy. Besides, our automated comparative analysis reveals that minipigs and mice exhibit the closest angular similarity to humans, reinforcing their value in comprehensive inner ear damage models. With the help of centerlines and landmarks, it is easy to automatically define the semicircular canal’s reference plane and calculate the angle parameters automatically.

However, there are some potential limitations and research directions that should be taken into consideration. Firstly, the fundamental issue about reconstruction is image resolution and could also be affected by the field of view, object size, and orientation^[8]^, factors that remain critical for refining NIHL model screening in the future. Because of the relatively lower resolution, supplementary modification is necessary for the procedure of reconstruction, which might overestimate the volume parameters by incorporating vessels, fluid, and other soft tissues in the gaps ^[8]^. Secondly, to propose an automatic measuring method suitable for different animals, only three animals (6 samples) in each species were included in this article. Interspecific comparisons and bilateral variations should be taken into analysis. Moreover, morphological changes like shrinkage induced by fixation, decalcification, and dehydration ^[8, 29]^, not necessary to micro-CT scan, were avoided in our preparation procedures, influencing the image quality. Based on the higher resolution of imaging, further reconstructions and measurements will be focused on the detailed anatomical inner ear structure such as the vestibular aqueduct and endolymphatic sac, which are pivotal in the pathogenesis of both congenital and noise-induced hearing loss.

In summary, this study presents an automated, centerline-based quantification method for the mammalian inner ear. Our data demonstrate that among common laboratory animals, the minipig possesses the highest morphological fidelity to the human inner ear across length, volume, and angular metrics. By providing a robust framework for inter-species comparison, this method facilitates the selection of superior animal models for NIHL, potentially accelerating the translation of therapeutic interventions for hearing disorders.

## 5. Conclusion

In conclusion, this study successfully established an automated 3D quantification framework for mammalian inner ear characterization using a micro-CT-based centerline method. Our comparative morphometric analysis demonstrates that miniature pigs exhibit the highest degree of anatomical and geometric similarity to the human inner ear among common experimental species, significantly surpassing traditional rodent models. These findings provide a robust mathematical foundation and morphological evidence for selecting miniature pigs as an ideal large animal model for NIHL research.

## Ethics Approval

All animal experiments were approved by the the Ethics Committee of Chinese PLA General Hosptial, approval **No. S2020-465-01** and performed in accordance with the ARRIVE guidelines and national regulations.

## Availability of Data and Materials

All relevant data are included within the article. Any additional materials (raw recordings, analysis scripts, or large imaging datasets) are available from the corresponding author upon reasonable request and in accordance with the journal’s data-sharing policy.

## Consent to Participate

Not applicable

## Consent for Publication

Not applicable

## Informed Consent

Not applicable (animal study).

